# 3D Chemotaxis Chip for Investigating Natural Killer Cell Migration Mechanisms

**DOI:** 10.1101/2025.04.21.649825

**Authors:** Madison N Temples, Suzanne Lightsey, Tiffany Conklin, Edward A Phelps, Blanka Sharma

## Abstract

**Introduction:** Natural killer (NK) cells are a promising tool for cancer immunotherapy, as they can rapidly recognize and kill cancer cells without prior knowledge of tumor-specific antigens while leaving healthy cells unharmed. However, a major challenge in NK cell-based therapies is their inadequate infiltration and function within solid tumors. Advancements in NK cell therapies for solid malignancies require a better understanding of the various factors that influence NK cell migration to and within the tumor microenvironment.

**Methods:** The objective of this study was to develop a chemotaxis chip with a tunable 3D hydrogel that enables the spatiotemporal analysis of NK cell migration.

**Results/Discussion:** By manipulating the 3D hydrogel or inhibiting key integrin and protease interactions, we found that NK cells heavily relied on protease-dependent migration but could leverage other mechanisms for faster migration. Additionally, when hyaluronic acid, an important extracellular matrix component in tumors, was incorporated into the hydrogel, NK cells migrated faster and farther in the chemotaxis chip.

**Conclusion:** This study establishes a novel migration assay to observe NK cell behavior in real-time, providing a platform for investigating the mechanisms of NK cell migration and identifying strategies to improve NK cell trafficking within solid tumors.

## 1. Introduction

Natural killer (NK) cells are potent effector cells of the innate immune system that eliminate cancer cells in an antigen-independent manner. Their mechanism of cancer cell recognition allows for the use of allogenic sources without the risk of graft-versus-host disease. As such, the adoptive transfer of NK cells has emerged as a promising therapy for cancer patients that bypasses some of the challenges associated with T cell therapies. NK cell therapy has induced remission or halted tumor progression in a subset of hematological cancer patients^1–5^; however, the outcome for patients with solid tumors has been less promising.^6–8^ Within solid tumors, NK cell infiltration is limited, and the cells that do infiltrate often remain in the tumor stroma^9–11^, failing to establish the cancer cell-NK cell contact necessary to exert their cytotoxic function. Indeed, clinical statistics suggest that the abundance of NK cells in the solid tumor microenvironment predicts outcomes in patients with several types of cancer, including melanoma^12–14^, breast cancer^15^, non-small cell lung cancer, pulmonary adenocarcinoma^16,17^, and gastric cancer^18^. Therefore, effective NK cell therapies that result in better infiltration and localization within the tumor are needed.

To reach a solid tumor, NK cells must first extravasate from the blood and traverse through the extracellular matrix (ECM) before exerting their cytotoxic functions.^19^ Recognition of chemokines secreted by tumors is critical for NK cell extravasation from the bloodstream and infiltration into the solid tumor.^20^ This is supported by the increased infiltration of engineered chemokine receptor-expressing NK cells into tumors^21–23^, and the greater presence of NK cells in tumors that contain engineered chemokine-expressing cancer cells.^24^ However, NK cell infiltration cannot be dependent on chemokine secretion alone, as even with high levels of tumor-secreted chemokines, endogenous NK cell^25^ and T cell^26^ infiltration can be limited. Indeed, immune cells also rely on interactions with extracellular substrates to reach their destination, but the extent or conditions under which the ECM acts as a substrate or a barrier to NK cell infiltration is unclear. In a syngeneic rat liver tumor model, adoptively transferred NK cells infiltrated into tumor models that lacked containment structures, comprised of collagen IV and laminin, but could not infiltrate tumor nodules that had containment structures.^27^ Conversely, the presence of vitronectin in the ECM of hepatic tumors corresponded to a greater presence and retention of leukocytes^28^, demonstrating the importance of integrin binding to the ECM for tumor retention. Given the complexity of the ECM in solid tumors, it remains challenging to determine whether components of the ECM promote or hinder NK cell migration.

To understand the role of the tumor ECM on NK cell infiltration, NK cell migration mechanisms in tumor-relevant ECMs need further study. The two major modes of 3D cell migration are mesenchymal and amoeboid^29^, with further classification in each category.^30^ Generally, the modes of migration are classified by their relative adhesion to the matrix and actin-myosin contractility. Cells using mesenchymal migration rely on actin polymerization at the leading edge to generate lamellipodia, and use integrins and associated focal adhesion proteins to interact with basement membranes.^29,30^ Further, mesenchymal migration is largely dependent on matrix metalloproteinases (MMPs) to degrade the ECM.^31^ Conversely, amoeboid migration, which resembles the movement of protist amoeba, is conventionally MMP-independent, but rather driven by actin protrusion of hydrostatic membrane blebs.^29,30^ Amoeboid migration requires high activity of Ras homolog gene family member A (RhoA) and rho-associated protein kinase (ROCK) for strong actin-myosin contractility to physically deform their cell body and squeeze through the surrounding pericellular matrix.^29^ While the mechanisms of migration for macrophages^32^, dendritic cells^33^, and T cells^34,35^ have been evaluated in three-dimensional (3D) systems, much less is known about NK cell migration in 3D. Understanding NK cell migration and employing rational design to improve their ability to navigate and sample the entire tumor could improve therapeutic efficacy against solid tumors.

Conventional methods for evaluating human NK cell migration rely on 2D culture, which lacks the 3D structure and microenvironment of solid tumors. Although these studies have provided insight into migration differences between resting and activated NK cells^36,37^ as well as the effect of surface topography on NK cell migration parameters^38^, their findings should be interpreted with caution, as cell migration mechanisms in 3D largely differ from 2D environments.^33^ To better mimic the tumor ECM, 3D systems made from natural materials like collagen^36,39,40^ and Matrigel^41^ have been used to study NK cell migration. In 3D collagen matrices, primary NK cells exhibit heterogeneous migration patterns^36,40^, with some cells moving slowly while others covered large distances at higher speeds.^36^ However, natural biomaterials are limited in their use for fundamental research due to their complex, variable, and ill-defined composition.^42,43^ Synthetic polymers, in contrast, offer more control over the physical, mechanical, and biological properties of the matrix. Poly(ethylene glycol) (PEG) is one of the most studied synthetic polymers and can be modified with diverse functional groups. PEG hydrogels are often formed via photopolymerization, a non-toxic technique that supports cell encapsulation. Our previous work showed that NK92 cells, a clinically relevant NK cell line, migrate into PEG-based hydrogels using MMPs and integrins toward a chemotactic point source.^44^ While these studies have advanced our knowledge of NK cell migration in 3D, they do not support real-time monitoring of migration in a tunable matrix. Therefore, there is a need for engineered platforms that integrate disease-relevant ECM architecture and enable temporal analysis of cell motility to identify key ECM components involved in NK cell migration.

In this work, we present a chemotaxis chip to study NK cell migration mechanisms in a PEG-based 3D hydrogel where the biochemical properties can be manipulated independently of the mechanical properties. Using the chemotaxis chip, we could quantify NK cell migration changes in response to different hydrogel compositions. Altogether, our study established a migration assay to study NK cell migration in real-time, which has further applications for understanding the mechanisms of migration and identifying novel therapeutic targets to enhance NK cell trafficking within the solid tumor.

## 2. Results and Discussion

### 2.1 Development of a Chemotaxis Chip for 3D NK Cell Migration

To create a 3D migration assay, we modified the commercially available µ-Slide Chemotaxis, a microfluidic tool amenable to time-lapse microscopy, to include a 3D PEG hydrogel in the observation area (Fig 1A). Briefly, the µ-Slide Chemotaxis features three separate chambers, with each chamber containing two symmetrical reservoirs bridged by a narrow observation channel. The observation chamber was filled with the desired PEG precursor solution (Fig 1B), polymerized using UVA light, and the reservoirs were filled with medium containing either primary NK cells or the chemoattractant. This setup allows for time-lapse imaging of NK cells migrating through the hydrogel.

**Figure 1.**
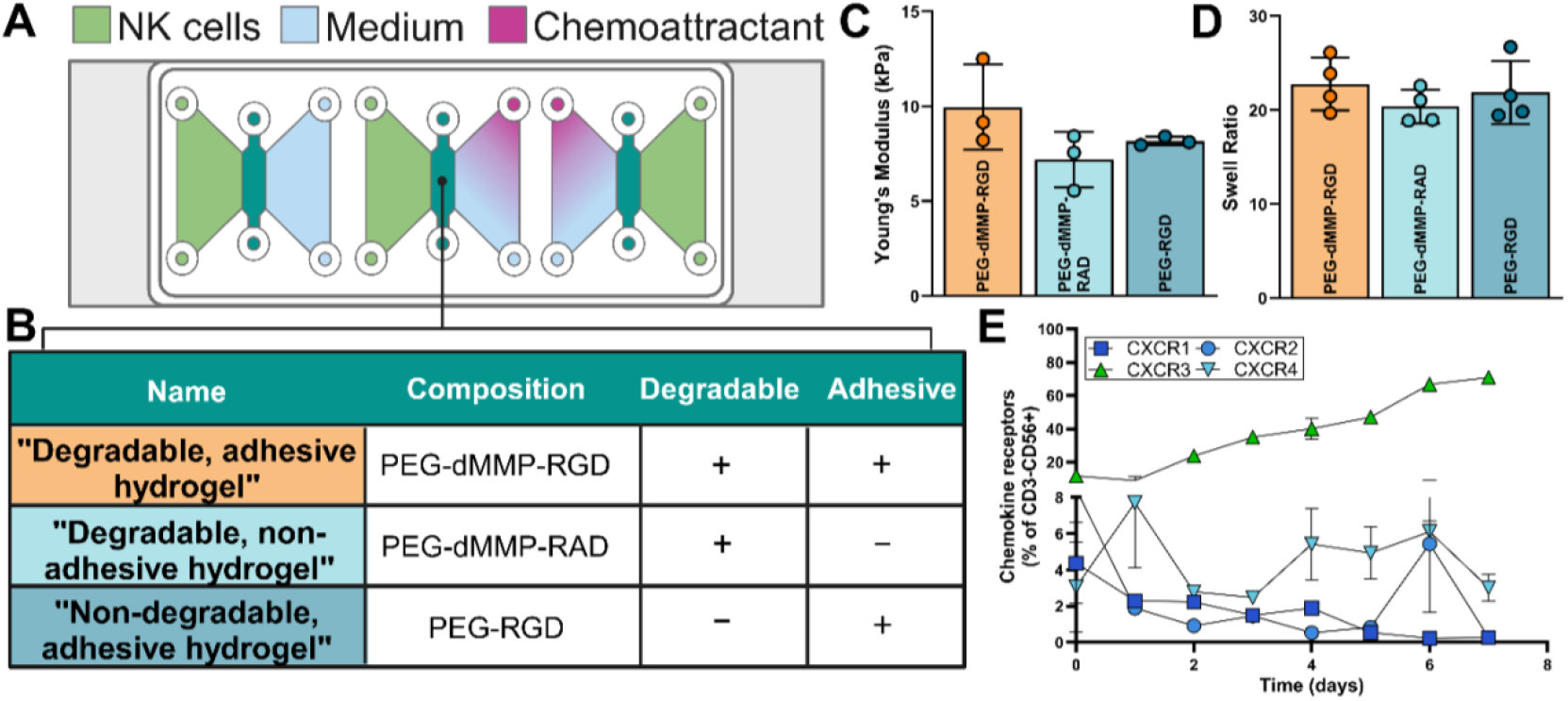
Establishment of the chemotaxis migration chip. (A) Schematic detailing an Ibidi chemotaxis µ-Slide chip set-up, with a hydrogel in the middle observation chamber. (B) Engineered PEG-based hydrogels and their properties which were crosslinked in the middle chamber of the chemotaxis chip. (C) Characterization of the stiffness of engineered hydrogels. (D) Quantification of the swell ratio of engineered hydrogels. (E) Characterization of primary NK cell chemokine receptor expression of CD3-CD56+ NK cells. Results are one donor representative of three donors.

To evaluate the mechanisms of NK cell migration, three PEG-based hydrogels with various degrees of adhesivity and degradability were engineered (Fig 1B). Using a synthetic material, we could alter the biochemical properties of the hydrogels without significantly changing the mechanical properties, as assessed by rheometry and hydrogel swelling properties (Fig 1C and 1D). The developed hydrogels have mechanical properties in the reported range for various solid tumors, which can range from 4-186 kPa depending on the tumor type and location.^45^ For example, breast cancer lesions have reported elastic moduli of 10-42 kPa, while healthy breasts have reported moduli of approximately 3.25 kPa.^46^ Although the system here is not specific to one tumor type, the Young’s modulus of 7-10 kPa is physiologically relevant and in line with those reported for many types of solid tumors.

Primary NK cells were isolated via negative selection from peripheral blood mononuclear cells of healthy human donors. Donor demographics are listed in Table S1. After isolation, the NK cells remained viable, were free of contaminating T cells, and had a high purity of CD3^-^CD56^+^ cells (Fig S1). To determine the appropriate chemokine to evaluate NK cell migration, we explored the expression of chemokine receptors on the isolated NK cells. There was a greater population of primary NK cells expressing CXCR3 after isolation and after 7 days of culture than the other chemokine receptors explored, CXCR1, CXCR2, and CXCR4 (Fig 1E). This finding is consistent with other studies reporting the predominant expression of CXCR3+ in primary NK cells compared to other chemokine receptors.^47,48^ The NK cells in this study were cultured in IL-2, which activates the cells and induces an upregulation of CXCR3 in NK cells relative to resting NK cells.^49^ Based on the expression of the chemokine receptors on the isolated NK cells, CXCL11 was selected as the chemokine for evaluating NK cell migration, as this is the most potent ligand for CXCR3.^50^

### 2.2 Characterization of the Concentration Gradient in the Chemotaxis Chip

The µ-Slide Chemotaxis is designed to maintain a stable, linear gradient; however, incorporating a hydrogel into the middle chamber may impact the stability and profile of the chemoattractant gradient. Therefore, it was necessary to characterize the gradient in the modified setup. To do so, we used 10 kDa FITC-labeled dextran, which has a comparable molecular weight to CXCL11, to represent the diffusion of CXCL11 in chemotaxis chips containing degradable, adhesive hydrogels. The gradient diffusion was evaluated over 24 hours and verified by fluorescent microscopy (Fig 2A). Within 12 hours, signal from the FITC-dextran was observed in the hydrogel. At 24 hours, the FITC-dextran signal stops increasing, indicating there is no longer a gradient across the observation area (Fig 2B). Since the chemokine gradient is lost after 24 hours, refreshing the chemokine daily was required to re-establish a chemokine gradient for long-term migration studies.

**Figure 2.**
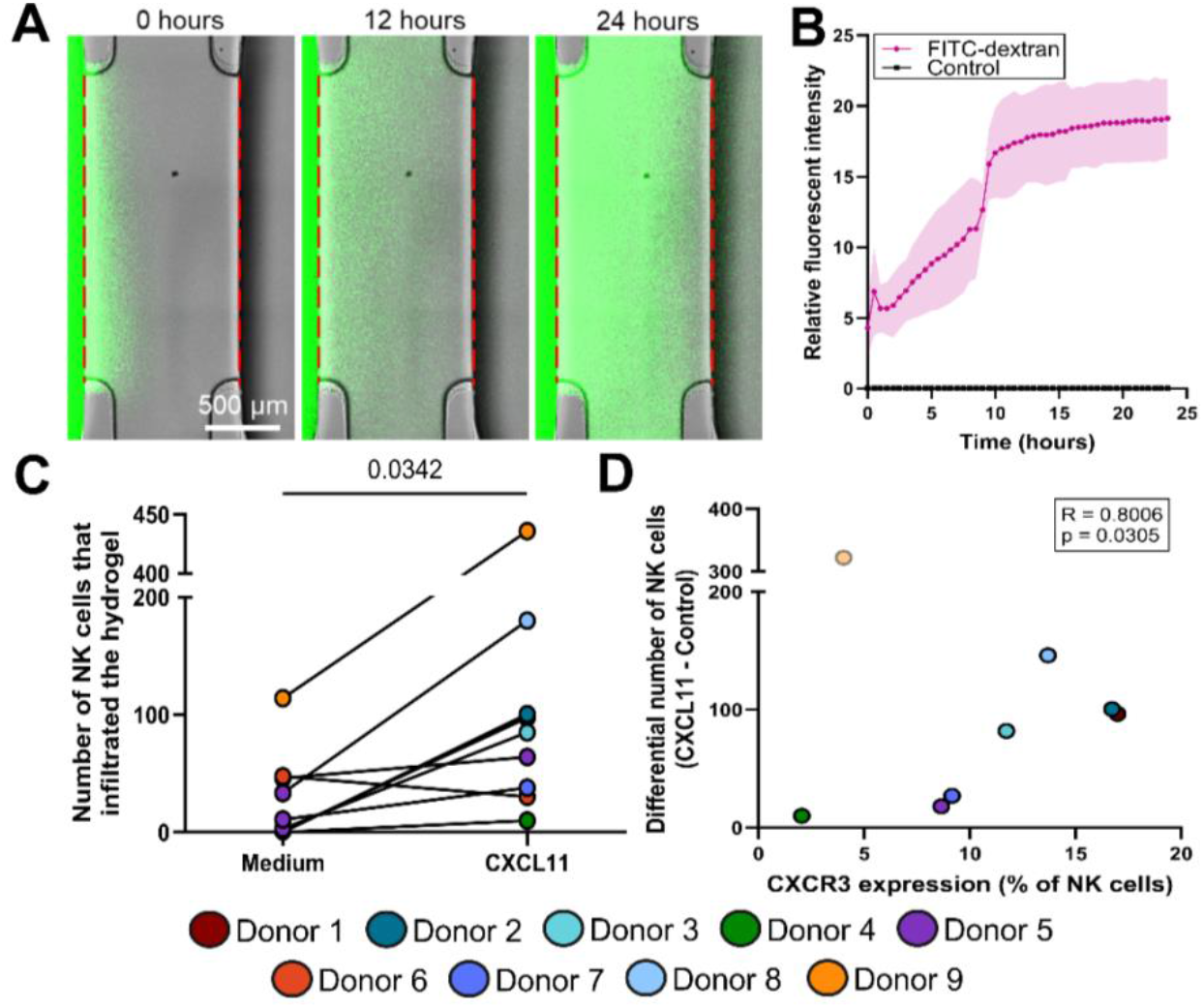
Directed NK cell migration into the 3D matrix in the chemotaxis chip. (A) Representative images of the diffusion of FITC-dextran (10 kDa) from the left chamber. The red dashed lines indicate the border of the observation area. (B) Quantification of the fluorescent intensity of the FITC-dextran gradient in the hydrogel relative to a control chamber that did not receive FITC-dextran. (C) Quantification of the average number of NK cells within the hydrogel after 5 days in culture in chambers with CXCL11 or medium only (control) (N=2-3 chambers per donor). (D) Correlation between the percent of CXCR3 expressing NK cells and the number of NK cells in the gel at day 5 in the CXCL11 chambers minus the NK cells in the medium (control) chambers. Of note, donor 6 was dropped from this analysis as this was the only donor to have an increase in the number of NK cells in the gel at day 5 in the medium (control) chamber compared to the CXCL11 chamber and donor 9 was treated as an outlier when calculating the Pearson correlation coefficient. Bars represent statistical significance.

### 2.3 NK Cell Migration Towards a Chemoattractant

Directed migration in the chemotaxis chip was evaluated by comparing NK cell migration towards CXCL11 or a media control. After primary cell isolation, the degradable, adhesive hydrogel was crosslinked in the observation area and rinsed in PBS; IL-2-containing media with CXCL11 was added to one reservoir and NK cells (4×10^6^ cells/mL) in IL-2-containing media were added to the opposite reservoir. The media and CXCL11 were refreshed daily, except in the control condition which only received media daily. After 2 days in the CXCL11-containing chemotaxis chip, NK cells were present in the hydrogel and cell numbers continued to increase during time in culture. CXCL11 was necessary for the cells to migrate into the hydrogel, as there were more NK cells in the hydrogels with CXCL11 than in the media control (Fig 2C). As expected, the donors who had greater expression of CXCR3 had more NK cells in the gel (Fig 2D). In this system, individual NK cells can migrate into the hydrogel and be tracked over time to evaluate migratory parameters (Video 1 and 2).

### 2.4 NK Cells’ Reliance on Mesenchymal Migration in 3D

Cells migrating in a mesenchymal manner largely use proteases to degrade ECM and integrins for adherence. As such, to determine NK cells’ reliance on mesenchymal mechanisms of migration, we evaluated their migration in hydrogels lacking either enzymatically degradable sites (PEG-RGD) or adhesion sites (PEG-dMMP-RAD). In both conditions, there was a decrease in the total number of NK cells that infiltrated into the hydrogel relative to the control hydrogel containing adhesion and degradable sites, indicating that matrix adhesion and degradation are necessary to initiate NK cell migration through the hydrogel (Fig 3A and 3E). Next, we imaged the observation chambers of the chemotaxis chip for 2 hours with a Leica TCS SP8 confocal microscope, acquiring images every 2 minutes, and quantified the dynamic movement of cells within these matrices. Three key metrics were used to analyze NK cell movement: (1) velocity, which refers to the rate of change of the cell’s position over time, (2) accumulated distance, which measures the total path length traveled by a cell over a given period, regardless of direction, and (3) Euclidean distance, which is the straight-line distance between two points. Since no cells infiltrated the non-degradable hydrogel, further migration parameters were not evaluated. When quantifying the migration parameters of NK cells within non-adhesive hydrogels, we found that NK cells migrated faster (1.4-fold) and traveled further, with a 1.3-fold increase in accumulated distance and a 1.2-fold increase in Euclidean distance, compared to cells in the adhesive hydrogel (Fig 3B-D). This data suggests that while NK cells rely on proteases to degrade the hydrogel, the presence of adhesion sites may slow their migration.

**Figure 3.**
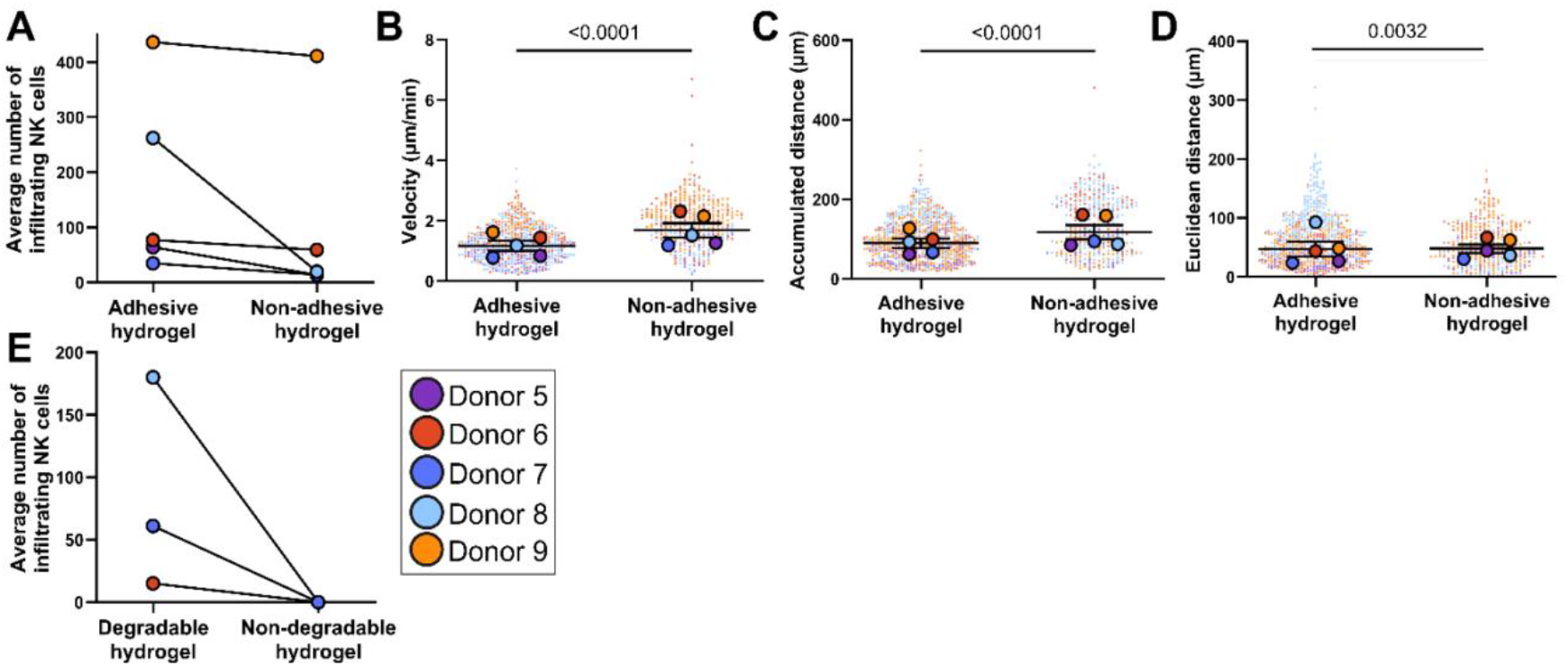
Evaluation of integrin and protease dependence of NK cells within 3D matrices. (A) Quantification of the average number of NK cells within the hydrogel after 5 days in culture in chambers with CXCL11 and either adhesive or non-adhesive hydrogels (N=2-3 chambers per donor). Quantification of individual cells migration parameters, including (B) velocity (μm/min), (C) accumulated distance (μm), and (D) Euclidean distance (μm). (E) Quantification of the average number of NK cells within the hydrogel after 5 days in culture in chambers with CXCL11 and either degradable or non-degradable hydrogels (N=2-3 chambers per donor). Data are presented as scatter dot plots with error bars representing SEM where large circles represent the donor’s average and small circles represent an individual cell. Bars represent statistical significance.

To further investigate the protease dependence of NK cell migration, we evaluated the migration of NK cells in the degradable, adhesive hydrogels in the presence of GM6001, a broad-range inhibitor of MMP. The cells were treated for two hours before and during the migration study with 5 μM of GM6001. For all donors evaluated, there were fewer GM6001-treated NK cells in the hydrogel relative to control NK cells, although in a non-significant manner (Fig 4A). However, the GM6001-treated NK cells that invaded the hydrogel migrated 23% faster than control NK cells (Fig 4B-D). Though we used GM6001 concentrations reported in literature^44^, it is conceivable that MMPs were not completely inhibited. Alternatively, enzymes other than MMPs may play a role in degrading the hydrogel, thereby allowing subsequent infiltration into the hydrogel. While the incorporated MMP degradable sequence used in this hydrogel was designed to be specific for a broad range of MMPs, the site has also been found to be susceptible to degradation by plasmin.^51^ Nonetheless, the data suggests that the MMP inhibitor may reduce the overall infiltration of the NK cells into the hydrogel; however, it does not completely block migration. NK cells that do infiltrate can migrate at similar or even faster velocities, perhaps by relying on alternative migration mechanisms, such as an amoeboid-like mode.

**Figure 4.**
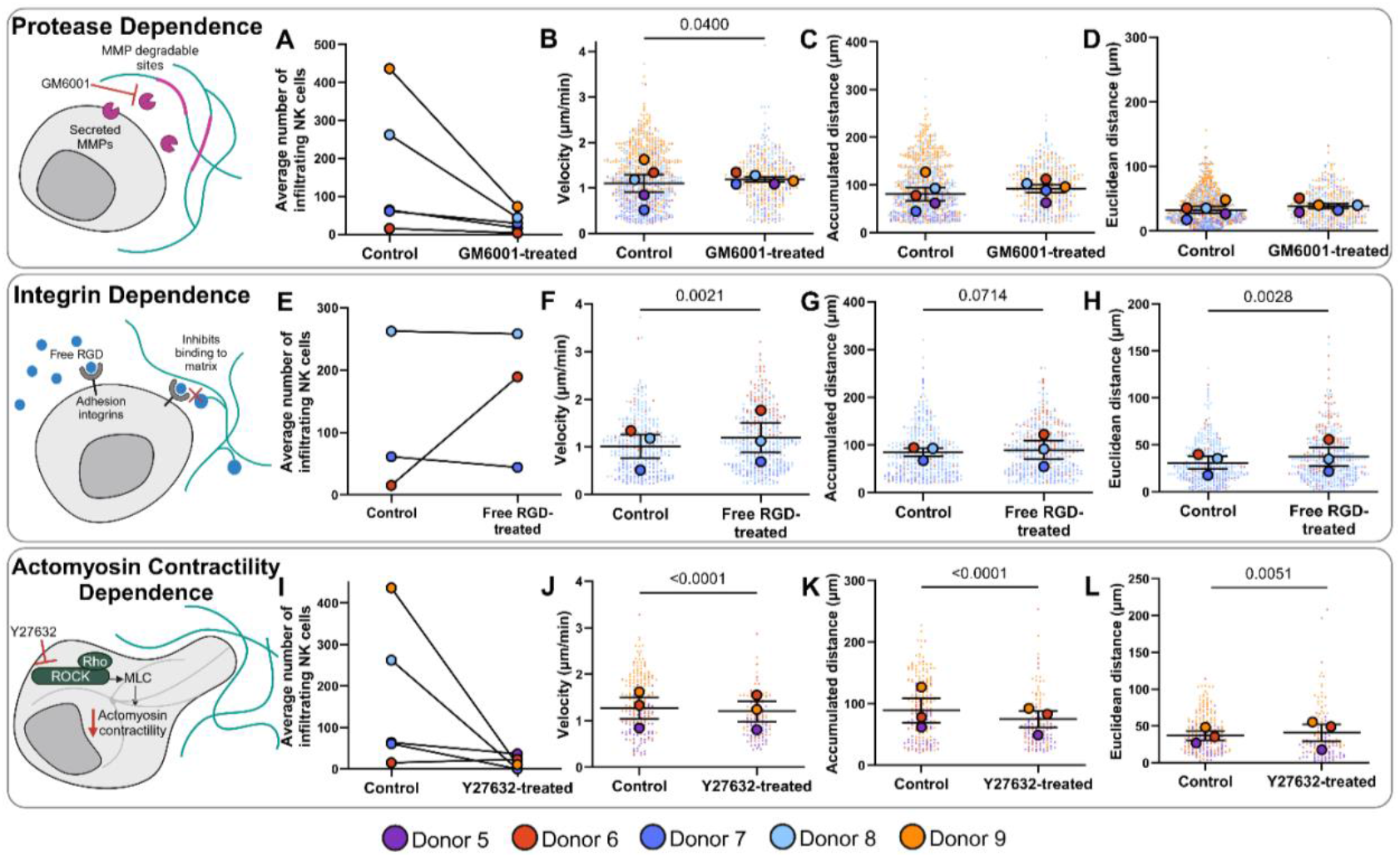
Characterizing NK cells mode of migration. (A) Evaluation of MMP-dependence in NK cell migration via inhibiting MMPs with GM6001 and quantifying the average number of NK cells that infiltrated the degradable, adhesive hydrogel (N=2-3 chambers per donor). Quantification of individual cells migration parameters, including (B) velocity (μm/min), (C) accumulated distance (μm), and (D) Euclidean distance (μm). (E) Evaluation of integrin-dependence in NK cell migration via blocking integrin adhesion to PEG-RGD using soluble/free RGD and quantifying the average number of NK cells that infiltrated the degradable, adhesive hydrogel (N=2-3 chambers per donor). Quantification of individual cells migration parameters, including (F) velocity (μm/min), (G) accumulated distance (μm), and (H) Euclidean distance (μm). (I) Evaluation of ROCK-dependence in NK cell migration via inhibiting ROCK with Y27632 and quantifying the average number of NK cells that infiltrated the degradable, adhesive hydrogel (N=2-3 chambers per donor). Quantification of individual cells migration parameters, including (J) velocity (μm/min), (K) accumulated distance (μm), and (L) Euclidean distance (μm). Data are presented as scatter dot plots with error bars representing SEM where large circles represent the donor’s average and small circles represent an individual cell. Bars represent statistical significance.

The role of cell adhesion in NK cell migration within the hydrogel was further explored by blocking integrins on NK cells with a pre-treatment of soluble RGD. Primary NK cells were treated with 50 μM of soluble RGD for two hours before and during the migration study using adhesive, degradable hydrogels. The treated NK cells that migrated into the hydrogel traveled 1.2 times faster and covered 1.2 times greater Euclidean distance than control NK cells (Fig 4F-H). Despite donor-to-donor variations, NK cells can migrate in either an integrin-dependent or -independent manner within this chemotaxis chip (Fig 4E). When NK cells utilized an integrin-independent mode of migration, they migrated faster and farther, consistent with an amoeboid-like mode of migration. The ability for integrin-independent 3D migration has also been observed in other immune cells. For example, while integrin adherence was necessary for 2D dendritic cell migration, it was not required for 3D migration in natural hydrogels.^52,53^ In contrast, T cells have been shown to physically adhere when migrating through a 3D collagen matrix, with contact guidance determining their forward migration and migratory turns, although no structural degradation or matrix remodeling occurred.^54^ The ECM composition, the availability of cell adhesion sites, and the presence of adhesion receptors on the cells likely influence the migration mechanisms of NK cells. Similarly, macrophages can switch their mode of migration depending on ECM characteristics.^55^ These findings suggest that NK cells are capable of adapting their migration strategies based on the ECM environment, potentially offering new insights into the regulation of immune cell trafficking and tissue infiltration.

### 2.5 Donor-Dependent Variation in ROCK-Mediated NK Cell Migration

While proteases and integrins are largely involved in mesenchymal migration, amoeboid-like migration primarily relies on the dynamics of the actin cytoskeleton and the ability of cells to deform and squeeze through small spaces without the need for protease activity or strong integrin-mediated adhesion. To explore the role of amoeboid migration in NK cell migration and invasion, Rho-associated coiled-coil kinase (ROCK) was inhibited with Y27632. ROCK regulates the cytoskeleton, particularly actin filaments, which are crucial for the blebbing characteristic of amoeboid cell migration. Primary NK cells were treated with 20 μM of Y27632 for two hours before and during the migration study in adhesive, degradable hydrogels. Overall, fewer Y27632-treated NK cells invaded the hydrogel, and those that did migrate moved more slowly and traveled shorter distances than untreated NK cells (Fig 4I-L). The dynamic parameters for cell migration were only evaluated for donors 5,6 and 9, as no treated cells were observed in the hydrogel for donors 7 and 8. Interestingly, there are striking donor-to-donor differences in the dependence on ROCK for migration within the chemotaxis chip. While donors 5 and 6 did not heavily rely on ROCK, donors 7-9 were particularly sensitive to ROCK inhibition with Y27632 (Fig 4I). These findings suggest donor variability in NK cell migration, with some subsets of NK cells possibly more adept at certain modes of migration. Further research is needed to identify and characterize these NK cell subsets and better understand the factors that govern their migration plasticity.

### 2.6 Hyaluronic Acid Enhances NK Cell Migration

To further demonstrate the application of the chemotaxis chip, we incorporated tumor-relevant ECM components and explored their role in NK cell migration. Solid tumors have higher hyaluronic acid (HA) content than healthy tissues^56,57^, which is associated with tumor growth and aggressiveness. Tumors contain both high molecular weight (HMW) and low molecular weight (LMW) HA, which result from the degradation of HMW HA by hyaluronidase produced by cancer and stromal cells.^57^ HMW HA has anti-inflammatory, anti-proliferative, and anti-angiogenic effects^58^, inhibiting tumor growth^59^ and reducing the production of MMPs^60^, while LMW HA promotes inflammation, proliferation, angiogenesis^58^, and metastasis.^61^ Interestingly, LMW HA can also reduce signaling pathways related to cancer cell behaviors.^58^ Immune cells also interact with HA, and it has been shown to enhance T cell migration, yet the influence of HA on NK cell migration parameters has not been studied.^62,63^ Given the limited understanding of the influence of the different molecular weights of HA on NK cell migration, we set out to further investigate this relationship in the established chemotaxis chip.

HMW HA and LMW HA were chemically methacrylated (MeHA) and incorporated into the degradable, adhesive hydrogels without changing the mechanical or physical properties of the hydrogel, as determined by Young’s modulus and swelling ratios (Fig 5A and 5B). While the inclusion of MeHA in the hydrogel did not alter the total number of NK cells that infiltrated into the hydrogel within the chemotaxis chip on average (Fig 5C), NK cells migrated 1.8-fold faster and traveled 1.7- or 1.4-fold greater accumulated distance in hydrogels containing either LMW MeHA or HMW MeHA, respectively (Fig 5D-E). These enhanced migration parameters in HA-containing hydrogels align with previous studies on HA’s impact on immune cell function and migration. For instance, NK92 cells cultured in HA hydrogels demonstrated improved cell proliferation, viability, and migration compared to cells cultured in 2D.^64^ Similarly, increased T-cell migration was observed in 3D hydrogels containing HMW HA (>200 kDa) compared to hydrogels without HA.^65^

**Figure 5.**
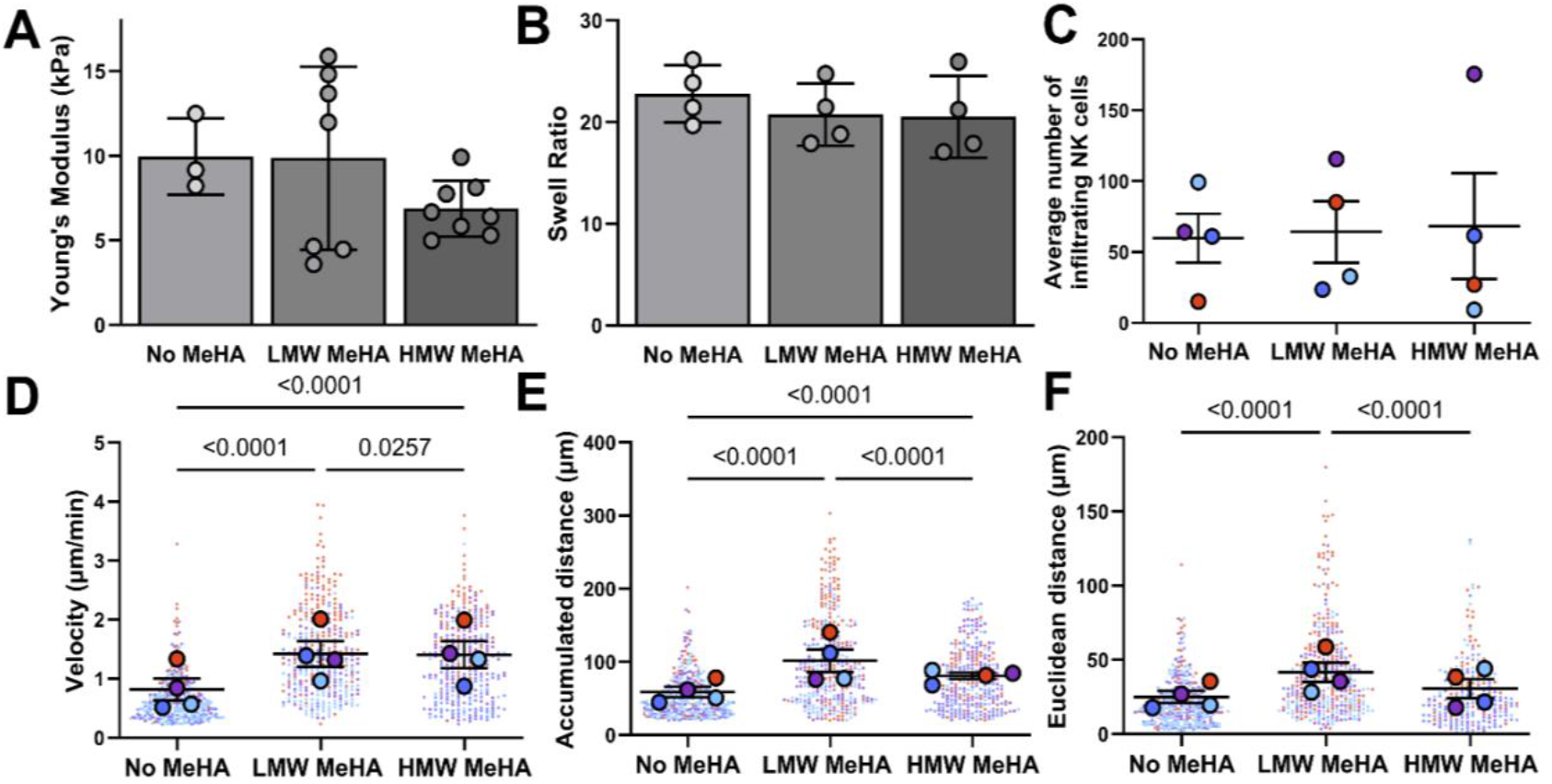
Effect of HA on NK cell migration. Either 500μg/mL of HMW MeHA, LMW MeHA, or no MeHA were added to degradable, adhesive hydrogels without significantly altering the (A) Young’s modulus or (B) swell ratio. (C) Quantification of the average number of NK cells within the hydrogel after 5 days in culture in chambers with CXCL11 and hydrogels containing MeHA. Quantification of individual cells migration parameters, including (D) velocity (μm/min), (E) accumulated distance (μm), and (F) Euclidean distance (μm). Data are presented as scatter dot plots with error bars representing SEM where large circles represent the donor’s average and small circles represent an individual cell. Bars represent statistical significance determined by one-way ANOVA followed by a Tukey’s multiple comparisons test.

Donor-specific differences were observed when comparing NK cell migration in hydrogels containing HMW MeHA or LMW MeHA. NK cells from donors 5-7 showed greater migration distance and speed in LMW MeHA, while NK cells from donor 8 migrated farther and faster in HMW MeHA. Since CD44 is the primary cell surface receptor for HA and plays a key role in migration and adhesion^66^, screening primary cells for CD44 expression levels may provide insights into the mechanisms governing their affinity for different molecular weights of HA.

## 3. Conclusion

NK cell migration is a vital process in cancer treatment, as it enables NK cells to effectively locate and penetrate solid tumors. Without proper migration, NK cells may fail to reach or fully sample the tumor, limiting their ability to target and eliminate cancer cells. Understanding and optimizing NK cell migration is therefore essential for improving their effectiveness in solid tumors, ensuring that these immune cells can efficiently engage and destroy cancerous tissue. The developed chemotaxis chip introduces a platform to investigate human NK cell migration mechanisms within an engineered 3D hydrogel, where the biochemical properties can be independently controlled. By modifying the commercially available µ-Slide Chemotaxis system, the chip enables the quantification of key NK cell migration parameters, such as cell infiltration, velocity, and migration distance, offering a multi-parametric readout of NK cell migration in a controlled 3D environment.

Our results demonstrate that NK cells rely heavily on protease-dependent mechanisms to penetrate the hydrogel but can utilize different modes of migration once within the hydrogel. Additionally, NK cell migration showed striking donor-to-donor variability in reliance on adhesion to RGD and/or ROCK-mediated migration, highlighting a personalized nature of NK cell migration. The inclusion of HA in hydrogels significantly increased both the velocity and distance traveled of NK cells across all donors. This suggests that HA may enhance NK cell infiltration into HA-rich tumors, which are common in cancers with high metastatic potential^67^, underscoring the importance of tumor ECM composition in NK cell trafficking.

While our platform addresses key aspects of NK cell migration mechanisms, it does not capture all the complexities of the solid tumor microenvironment. Instead, our goal was to strategically incorporate critical elements of the tumor milieu to better understand how specific variables, like integrins and proteases, impact NK cell migration. Although it is not feasible to integrate every variable of the tumor microenvironment into a single *in vitro* model, our system replicates critical physiological features to shed light on the impact of specific variables on NK cell migration. The findings align with similar research on immune cell migration, validating the accuracy of the chemotaxis chip. For instance, primary NK cells exhibited dynamic migratory behaviors, alternating between periods of arrest and movement^68^ and similar findings have been reported in other immune cells, such as macrophages^55^ and T cells^69^, where plasticity in migration modes was found to depend on the 3D environment. These observations emphasize the importance of studying immune cell migration in 3D models and highlight the chip’s value as a modular platform for further exploration.

Ultimately, we envision the chemotaxis chip as part of a broader network of *in vitro* models designed to capture different facets of immune cell behavior and migration. To fully understand the therapeutic potential of NK cells in solid tumors, additional assays evaluating NK cell cytotoxicity, degranulation, and cytokine release are necessary. These complementary assays will provide a more comprehensive understanding of NK cells with differing migration profiles and their potential as immunotherapy candidates for solid tumors. While the developed chemotaxis chip was used to study fundamental mechanisms of NK cell migration in solid tumors, it may also have broader applications in areas such as wound healing, transplant rejection, and placental development. In conclusion, the chemotaxis chip is a step toward developing tailored strategies to enhance NK cell therapies, improving their ability to infiltrate and act within solid tumors.

## 4. Methods

### 4.1 Materials

Leukocyte reduction chambers from healthy donors were purchased from LifeSouth Gainesville after receiving informed consent from all participants. Peripheral blood mononuclear cells (PBMCs) were isolated via Ficoll centrifugation. NK cells were isolated from the PBMCs via negative selection magnetic separation (Untouched NK cells Dynabeads, Invitrogen). NK cells were cultured in RPMI 1640 (without L-glutamine), 10% fetal bovine serum, 1% penicillin-streptomycin, 1% L-glutamine, 1% non-essential amino acids, 1% sodium pyruvate, MycoZap, and 100 U/µL IL-2 (Peprotech, Inc.). NK cells were kept at a density of 1×10^6^ cells/mL in culture for 7 days at 37°C and 5% CO_2_. The study was approved by the University of Florida Institutional Review Board (IRB202002072). The demographics of donors are listed in Table S1.

PEG-diacrylate (PEG-DA, MW 3.4 kDa) and acrylate-PEG-succinimidyl valerate ester (acryl-PEG-SVA, MW 3.4 kDa) were purchased from Laysan Bio Inc. The matrix metalloproteinase (MMP) degradable sequence, GGVPMS↓MRGGK, (MW 1076.31 Da) was purchased from Biomatrix. Cyclo(Arg-Gly-Asp-D-Phe-Lys) (RGD, MW 603.68 Da) and cyclo(Arg-Ala-Asp-D-Phe-Lys) (RAD, MW 617.71 Da) were purchased from Biosynth Inc. Dialysis tubing, Spectra/Por6 2000 MWCO, was purchased from Spectrum Laboratories. 2-Hydroxy-4’-(2-hydroxyethoxy)-2-methylpropiophenone (Irgacure 2959) was purchased from Spectrum Laboratories. All inhibitors were purchased from Calbiochem. All other materials were obtained from Thermo Fisher Scientific.

### 4.2 Fabrication of Chemotaxis Chip

The chemotaxis chip was modified from the commercially available microfluidic device (Ibidi µ-Slide Chemotaxis) to support the migration of primary NK cells through an engineered 3D hydrogel. The device consists of three individual chambers, each with two symmetrical reservoirs connected by a narrow observation channel. Our experimental design included one reservoir with NK cells, the other containing medium (either with or without CXCL11), and the observation channel containing the hydrogel, unless specified otherwise. Briefly, 5.12 µL of the hydrogel precursor solution was injected to the top port of the observation chamber with the ports plugged on the outer reservoirs as detailed by Ibidi. 6 µL of air was aspirated from the bottom port of the observation chamber to pull the precursor solution through. This process was repeated for all three chambers. The slide was then photopolymerized by the OmniCure S1000 light (Excelitas Technologies, Corp.) for 3 minutes using 4.0 mW/cm^2^ long wave ultraviolet A (UVA) light. For donors 5-7, the slide was photopolymerized by OmniCure S2000 light (Excelitas Technologies Corp.) for 2.5 minutes using 4.24 mW/cm^2^ long wave UVA light. The hydrogels were washed by removing the plugs from the reservoirs and filling the ports with 65 µL of PBS. Then the PBS was removed from one reservoir and 65 µL of NK cell culture medium was added via the top port. Then, the PBS was removed from the opposite reservoir and 75 µL of NK cells suspended in medium (at 4×10^6^ cells/mL) was added via the top port. Lastly, 10 µL of 1 ng/µL CXCL11 in medium or medium alone, for the control, was added to the top port of the reservoir containing media only. For studies quantifying the concentration gradient, FITC-dextran was used instead of CXCL11.

Primary NK cells were seeded in their respective reservoirs one or two days after isolation. Each chemotaxis chip would investigate one experimental parameter with the following set up: chamber 1 had NK cells in the left reservoir and medium only (control) in the right reservoir, chamber 2 had NK cells in the left reservoir and CXCL11 in the right reservoir, and chamber 3 had NK cells in the right reservoir and CXCL11 in the left reservoir (Fig 1A). Having the chemokine in different reservoirs accounts for any external tilt the chip may experience during culture or imaging.

### 4.3 Cell Culture in Chemotaxis Chip

The chemotaxis chip was cultured at 37°C and 5% CO_2._ While in culture, the media and the chemoattractant gradient were refreshed daily. The medium was removed from the individual CXCL11 or control reservoirs and 65 µL of NK cell culture medium was added to each of the CXCL11 or control reservoirs via the top port. Then, 75 µL of NK cell culture medium was added to the NK cell-containing reservoirs via the top port and approximately 60 µL of spent NK cell culture medium was removed from the bottom port. Lastly, 10 µL of CXCL11 in medium (1 ng/µL) or medium alone for the control group was added to the top port of the medium containing reservoirs. The chemotaxis chips were imaged using a Leica TCS SP8 confocal microscope (Leica Microsystems) on the 5^th^ day of culture, with the media and chemokine being refreshed prior to imaging.

### 4.4 NK Cell Inhibition

For studies involving inhibitors, NK cells were pre-treated for 2 hours with the inhibitor and cultured in inhibitor-containing medium for the duration of the experiment. The inhibitors did not affect NK cell viability (Fig S2). In all inhibitor studies, the degradable, adhesive hydrogel (PEG-dMMP-RGD) was used. To inhibit MMPs, NK cells were treated with 5 µM of the broad-range MMP inhibitor GM6001. To inhibit integrin binding, NK cells were treated with 50 µM of a soluble RGD peptide. To inhibit ROCK, NK cells were treated with 20 µM of Y27632.

### 4.5 Peptide Functionalization and Polymer Synthesis

The MMP-sensitive peptide sequence VPMS↓MRGG was included in the hydrogel to support cell migration. The peptide was modified to be capped on either end of a PEG arylate group upon conjunction as previously described.^70–74^ Briefly, the terminal and lysine amine groups of the MMP degradable sequence were reacted with the SVA of an acryl-PEG-SVA in 50 mM sodium bicarbonate (pH = 8.0) at a 1:2 molar ratio for 4 hours. The reaction product was dialyzed for 24 hours against water to remove unreacted reagents, lyophilized for 48 hours and stored at -20°C until use.

Cell adhesion was supported by immobilizing cell adhesion ligands to the crosslinked network. RGD functionalized PEG, acryl-PEG-RGD (PEG-RGD, MW ∽4005 Da), which mimics the adhesion site on vitronectin^75–77^, was synthesized as previously described.^72–74^ Briefly, the terminal amine of cyclo(Arg-Gly-Asp-D-Phe-Lys) (cRGD) reacted with acryl-PEG-SVA in 50 mM sodium bicarbonate (pH = 8.0) at a 1.05:1 molar ratio for 4 hours. The reaction product was dialyzed for 24 hours against water to removed unreacted reagents, lyophilized for 48 hours, and stored at -20°C until use. Acryl-PEG-RAD (PEG-RAD, MW ∽4013 Da), a control with no integrin binding site was synthesized in the same manner, by reacting the terminal amine of cyclo(Arg-Ala-Asp-D-Phe-Lys) with the SVA of an acryl-PEG-SVA.

Methacrylated hyaluronic acid (MeHA) was synthesized as previously described.^78^ Briefly, hyaluronic acid (HA) was dissolved in a 50:50 ratio of PBS and acetone solution at 10 mg/mL overnight. Triethylamine was added and stirred for 30 minutes, followed by the addition of glycidyl methacrylate and stirred overnight. The stoichiometric ratio of glycidyl methacrylate added to each HA monomer was 20X. MeHA solution was precipitated by pouring the dissolved solution into a large beaker of acetone and stirring with a glass rod to collect the precipitate. Once the solution precipitated, MeHA was rinsed with fresh acetone and dissolved in PBS overnight, followed by another precipitation in acetone. MeHA solution was then dialyzed for 72 hours against PBS, followed by another dialysis for 36 hours against water. MeHA solution was sterile filtered, flash frozen, lyophilized for 5 days, and stored at -20°C until use. This process was done for high molecular weight HA (750-1000 kDa, bacteria-derived, Lifecore Biomedical) and low molecular weight HA (10-30 kDa, bacteria-derived, Sigma Aldrich).^57^

### 4.6 Peptide Functionalization Analysis

The PerkinElmer Frontier spectrometer was utilized to obtain the attenuated total reflectance Fourier transform infrared spectra (ATR-FTIR) of lyophilized PEG-dMMP-PEG, PEG-RGD, and PEG-RAD. The spectrometer was equipped with desiccate Ga-coated KBr optics, a temperature-stabilized deuterated triglycine sulfate detector, and one reflection diamond/ZnSe universal ATR sampling accessory.^79^ ATR-FTIR selected characteristic bands for PEG-SVA: C-H stretching 2880 cm^-1^ and C-O stretching 1102 cm^-1^. ATR-FTIR selected characteristic bands for cRGD peptide: aromatic ring 1644 cm^-1^. ATR-FTIR selected characteristic bands for dMMP peptide: C=O stretching 1645 cm^-1^ and C-S stretching 660 cm^-1^. Confirmation of PEG functionalized for PEG-dMMP-PEG and PEG-RGD are shown in Figures S3 and S4, respectively.

The degree of methacrylation of HMW and LMW MeHA was calculated via proton nuclear magnetic resonance (HNMR) spectroscopy. HNMR spectra were obtained using a 600 MHz Bruker Avance III spectrophotometer. Deuterium oxide (ACROS Organics) was used as a solvent and analysis was done at room temperature. The relative peak integration was performed between methacrylate protons (peak at 1.8 ppm) and methyl protons already present on the HA backbone (peak at 1.9 ppm), as previously described.^78^ This is referenced as the “degree of methacrylation” to mean the percentage of HA monomers containing methacrylated groups. For HMW and LMW MeHA, the degree of methacrylation was 21.27% and 19.97% (Fig S5).

### 4.7 Characterization of Hydrogel Properties

The precursor solutions for the different hydrogel formulations made in PBS, are listed in Table S2. The photoinitiator, Irgacure 2959, was added to each precursor solution (0.05% w/v) before polymerization. Briefly, hydrogels were formed in cylindrical silicon molds (Grace Biolabs) 8 mm diameter by 1.7 mm depth. The precursor solution was added to the silicon mold and photopolymerized by the OmniCure S2000 light for 3 minutes at 4.24 mW/cm^2^ long wave UVA light. After polymerization, the hydrogels were incubated in PBS at 37°C and 5% CO_2_ for 24 hours. The viscoelastic properties were examined using an Anton Paar MCR 302 rheometer with parallel plate geometry according to established methods.^80,81^ An 8 mm sandblasted top load cell was used to decrease slipping during testing. For an approximation of physiological conditions, the Peltier plate was preheated to 37°C, and the samples were enclosed in a humidity chamber. For each hydrogel composition, amplitude sweeps from 0.01% to 100% strain were first conducted to determine the linear viscoelastic region (LVE). The storage (G’) and loss (G”) moduli were then determined from frequency sweeps from 0.1 to 100 Hz at the strain value in the middle of the LVE, chosen as 0.1% strain for all hydrogel compositions. For all hydrogel compositions, shear moduli (G*) were determined from equation 1 at 1 Hz and used to calculate the Young’s modulus (E) using equation 2 with a Poisson’s ratio (U) of 0.5.^82–87^ For each condition, three samples were used for the amplitude sweeps and four samples were used for the frequency sweeps, wherein each hydrogel was used only for one test.

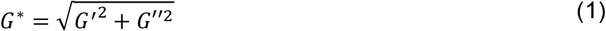

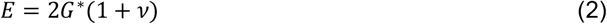

The swell ratios of the hydrogels were determined as previously described.^70^ Precursor solutions (10 mL) were photopolymerized as previously described. After polymerization, the hydrogels were incubated in PBS for 24 hours at 37°C and 5% CO_2_. The hydrogels were carefully blotted with a Kim wipe to remove excess surface liquid, and the swollen wet weight (*m*_*s*_) was measured. Hydrogels were lyophilized for 24 hours and the dry weight of that hydrogel (*m*_*d*_) was measured. The swell ratio at 24 hours was calculated by equation 3.

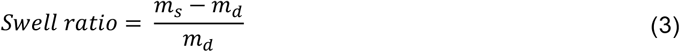

Further, for each composition, the weight fractions of the gel after crosslinking (q_F_, 0 hours) and when swollen (q_w_, 24 hours) were determined by equations 4 and 5, respectively. Here, *m*_*c*_is the wet weight of the hydrogel immediately after crosslinking and *m*_*d*_ is the dry weight of that hydrogel, and *m*_*s*_ is the swollen gel wet weight and m_d_ is the dry weight of that hydrogel.

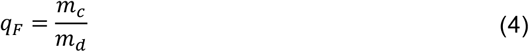

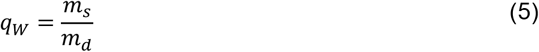

Additionally, it was necessary to calculate the volume fraction of the polymer of the swollen gel (*v*_2,*s*_) and the volume fraction of the polymer after crosslinking (*v*_2,*r*_) by equations 6 and 7. Here, the density of PEG (*ρ*_*p*_ = 1.12 *g*/*cm*^3^) and the density of the solvent, PBS (*ρ*_*sol*_ = 1.00 *g*/*cm*^3^) are needed for the calculations.

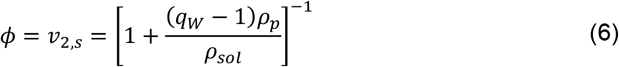

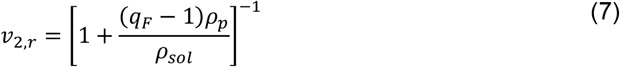

The molecular weight between crosslinks, M_c_, can be calculated by the Peppas-Merrill equation, shown in equation 8.^88–90^ The Peppas-Merrill equation is a modification of the Flory-Rehner equation and applies to swollen networks of crosslinked polymers prepared under conditions that the crosslinks were introduced in solution.^88–90^ Here, the number-average molecular weight of the polymer (*M*_*n*_) for PEG-dMMP-PEG was 7900 Da, the specific volume of the polymer 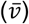 for PEG was 0.893 cm^3^/g,^91,92^ the molar volume of solvent (*V*_1_, for water) was 18 g/mol,^90^ and the polymer solvent thermodynamic interaction parameter (χ) for PEG and water was 0.43.^91–93^

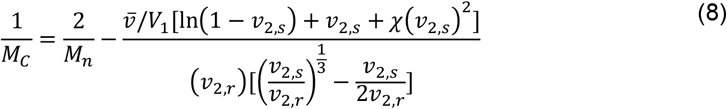

The mesh sizes of the hydrogels were then determined by equations 9 and 10.^89,94^ The average end-to-end distance of the polymer chain in the unperturbed state 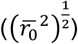 was determined by equation 4-9, where the molecular weight of the repeating unit (*M*_*r*_) for PEG was 44, the average bond length (*l*) for PEG was 0.154 nm^94^ and the Flory characteristic ratio of the polymer (*C*_*n*_) for PEG was 4.0.^91,92^

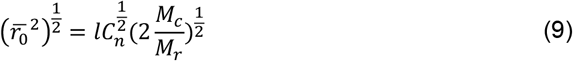

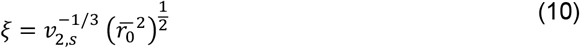

### 4.8 Flow Cytometry

Viability, purity, NK cell subset, and chemokine receptor expression were assessed by flow cytometry after isolation and the subsequent days in culture. Cell viability was assessed by live/dead^TM^ fixable Aqua Dead Cell Stain Kit (Invitrogen). NK cell purity and phenotype were characterized by staining with conjugated with conjugated mouse anti-human antibodies against CD3 (Clone SK7, BD Biosciences), CD56 (Clone B159, BD Biosciences), and CD16 (Clone 3G8, BD Biosciences). NK cells chemokine receptors were characterized by staining with conjugated mouse anti-human antibodies against CXCR1 (Clone 5A12, BD Biosciences), CXCR2 (Clone 6C6, BD Biosciences), CXCR3 (Clone 1C6/CXCR3, BD Biosciences), and CXCR4 (Clone 12G5, BD Biosciences). NK cells are defined as CD3- and CD56+ (representative gating strategy shown in Fig S6). A minimum number of 100,000 cells were analyzed using a BD FACS Celesta Flow Cytometer and Flowjo Software v10 (FLOWJO, LLC Data analysis software).

### 4.9 Confocal Microscopy and Image Analysis

Chemotaxis chips with primary NK cells were imaged using a Leica TCS SP8 confocal microscope with HC PL APO CS2 10x/0.4 Dry air objective. The microscope was pre-warmed to 37°C and pre-set to 5.5% CO_2_, the humidity chamber had a dish filled with synergy water and two Kim wipes rolled up and soaked with synergy water to prevent evaporation. Additionally, chemotaxis chips were taped down on all sides (avoiding the observation areas) so that the chip would not shift during imaging. NK cell migration in the chip was imaged with spatial temporal time-course imaging, which was acquired at 512 × 512-pixel resolution. Chemotaxis chips were imaged for 1.5-2 hours, with images being acquired every 2 minutes. Images were acquired with a pinhole of 11.31, a zoom of 4.00, and a z-stack size of 10-25 µm. FITC-dextran diffusion chips were imaged with spatial temporal time-course imaging, which was acquired at 512 × 512-pixel resolution. The chips were imaged for 24 hours, with images being acquired every 30 minutes. Images were collected with a pinhole of 11.31, a zoom of 1.25, and a z-stack size of 100 µm. For all studies, the z-stack images were projected onto a 2D image using volume intensity projections.

Image analysis was performed using Fiji (National Institute of Health).^95^ NK cells in the hydrogel at time 0, the first image of the series, were counted using the Cell Counter plugin. Then the cell tracks were manually tracked with the Manual Tracking plugin in Fiji (National Institute of Health) for the first 90 minutes of migration. The CSV file with x and y locations from the Manual Tracking plugin were then imported into a Wolfram Mathematica code developed to take those tracks and determine migration parameters. First, the cell tracks were adjusted so the (x, y) position was (0, 0) at 0 hour for each cell. From these modified cell tracks the accumulated distance, Euclidean distance, and velocity were evaluated for each cell. The accumulated distance is an average of all the cells movements measured in between all analyzed images (slices), m, calculated by equation 11, d_i,j_ is the displacement of the cell number I from image j-1 to image j. The Euclidean distance is an average of all the cells length of the individual cell start and end and calculated by equation 12. The velocity (V) is calculated by equation 13. Only tracks with an accumulated distance of 20 μm (2x the cell body) or more were included in the results.

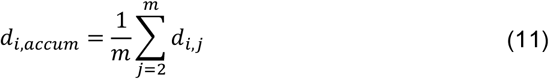

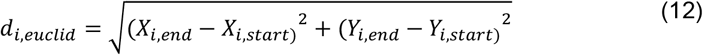

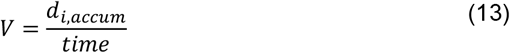

### 4.10 Statistical Analysis

GraphPad Prism V9 (GraphPad Software, San Diego, CA, USA) was used to perform statistical analyses. For flow cytometry data, a one-way ANOVA and post hoc analysis with Dunnett’s correction was performed to identify differences within one group, against day 0. Additionally, a one-way ANOVA and post hoc analysis with Tukey correction was performed to identify differences between groups on day 7 of culture. For the need for CXCL11 in the hydrogel, a paired Student t-test was performed. For all primary NK cell migration results, unpaired Student t-tests were performed to compare each experimental group to the control group. Statistical significance was considered as p ≤ 0.05. All numerical data are shown as donors, with the mean ± standard error of the mean (SEM) overlaid on the graph. Of note, not all experiments were evaluated on all 9 NK cell donors. For Figure 2, each point represents the results from one donor, with the CXCL11 results being an average of both chambers with the chemokine. For graphs of velocity, accumulated distance, and Euclidean distance each small point represents an individual cell, and large circles represent the donor’s average.

## Data availability

The Supplemental Material supporting this article has been included as part of the ESI.†

## Acknowledgements

A portion of this work was performed in the McKnight Brain Institute at the National High Magnetic Field Laboratory’s Advanced Magnetic Resonance Imaging and Spectroscopy (AMRIS) Facility, which is supported by National Science Foundation Cooperative Agreement No. DMR-1644779 and the State of Florida. This work was funded by the National Science Foundation (NSF) Division of Chemical, Bioengineering, Environmental and Transport Systems (CBET) Faculty Early Career Development (CAREER) Program (Award #1845728), Graduate Research Fellowship Program (GRFP) (Award #1000308740), and the Leo Claire & Robert Adenbaum Foundation. Research in the Phelps lab is supported by NIH grants R01 DK132387, R01 DK124267, and Breakthrough T1D grant 2-SRA-2023-1313-S-B. Figure 1 and 4 were created with Biorender.com.

