## Supplemental Material for "3D Chemotaxis Chip for Investigating Natural Killer Cell Migration Mechanisms"

---

| Donor number | Sex | Age | Ethnicity | ABO |
|---|---|---|---|---|
| 1 | Female | 53 | Caucasian | A- |
| 2 | Male | 34 | Caucasian | O+ |
| 3 | Male | 23 | Caucasian | A+ |
| 4 | Female | 24 | Hispanic | A- |
| 5 | Female | 62 | Caucasian | A+ |
| 6 | Female | 27 | Caucasian | O+ |
| 7 | Female | 33 | Caucasian | A+ |
| 8 | Male | 55 | Caucasian | A+ |
| 9 | Female | 70 | Caucasian | O+ |

**Table S1**. NK cell donor demographics.

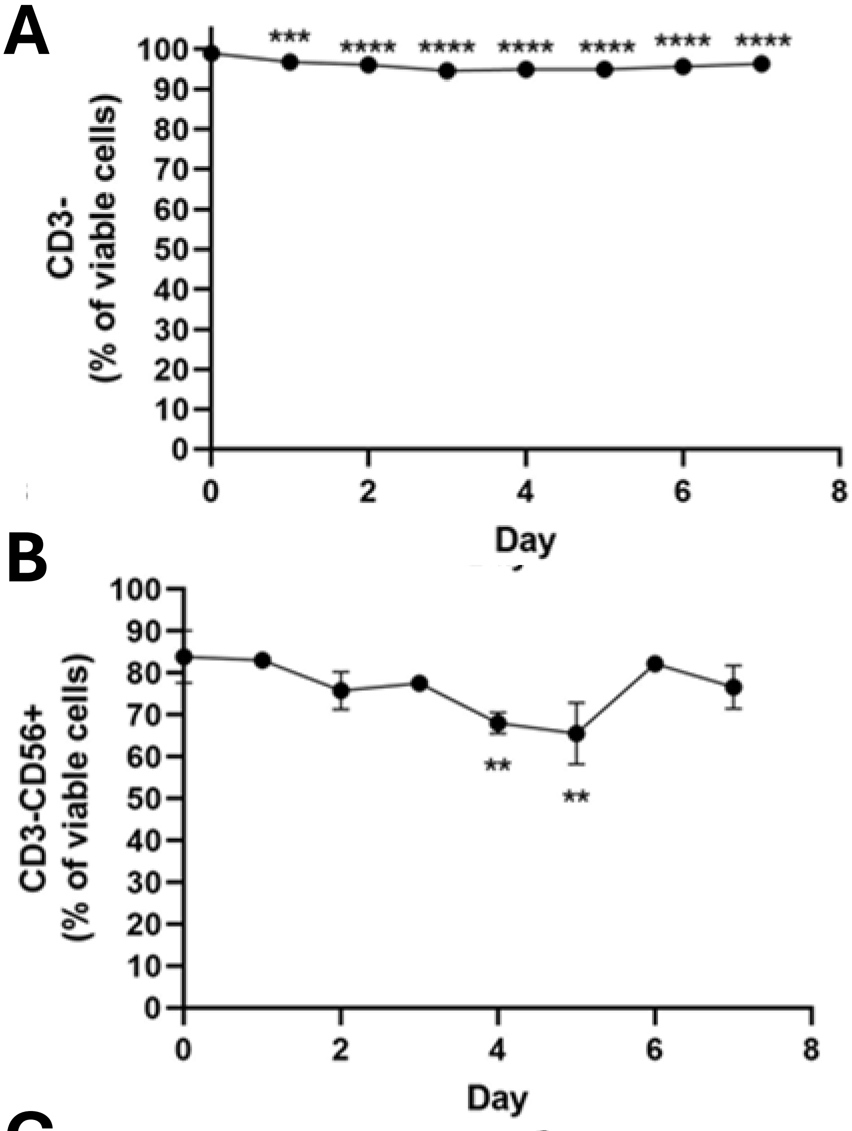

**Figure S1. Characterization of primary NK cells.** After primary cell isolation and subsequent days in culture, NK cells were characterized via flow cytometry. (A) Isolated NK cells were negative for CD3 and free of contaminating CD3+ T cells. (B) Purity of CD3-CD56+ NK cells or the isolated cells. (C) Characterization of the NK cell populations. Results are one donor representative of 3 donors. Statistical significance was determined using a Student's T-test; **p<0.01, ****p<0.0001

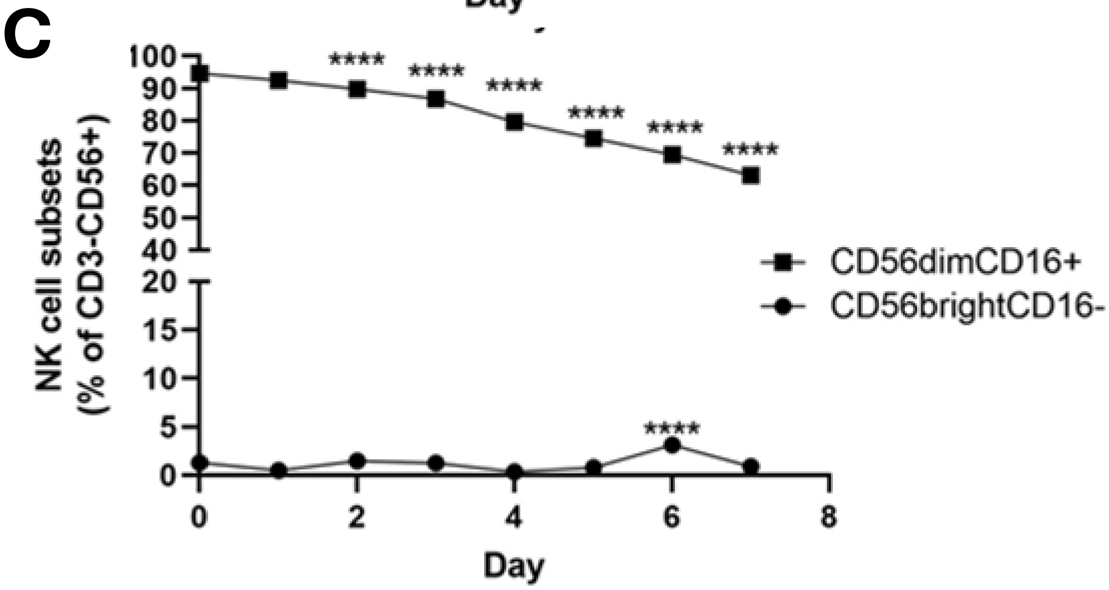

| Hydrogel name | Precursor Solution Composition |
|---|---|
| Degradable, adhesive hydrogel | 10% (w/v) PEG-dMMP-PEG (8 kDa) and 5 mM PEG-RGD |
| Non-degradable, adhesive hydrogel | 10% (w/v) PEG-PEG-PEG (10 kDa) and 5 mM PEG-RGD |
| Degradable, non-adhesive hydrogel | 10% (w/v) PEG-dMMP-PEG (8 kDa) and 5 mM PEG-RAD |
| LMW MeHA | 10% (w/v) PEG-dMMP-PEG (8 kDa), with 5 mM PEG-RGD, and 500 µg/mL LMW MeHA (10-30 kDa) |
| HMW MeHA | 10% (w/v) PEG-dMMP-PEG (8 kDa), with 5 mM PEG-RGD, and 500 µg/mL HMW MeHA (750-1000 kDa) |

**Table S2**. Hydrogel precursor solutions.

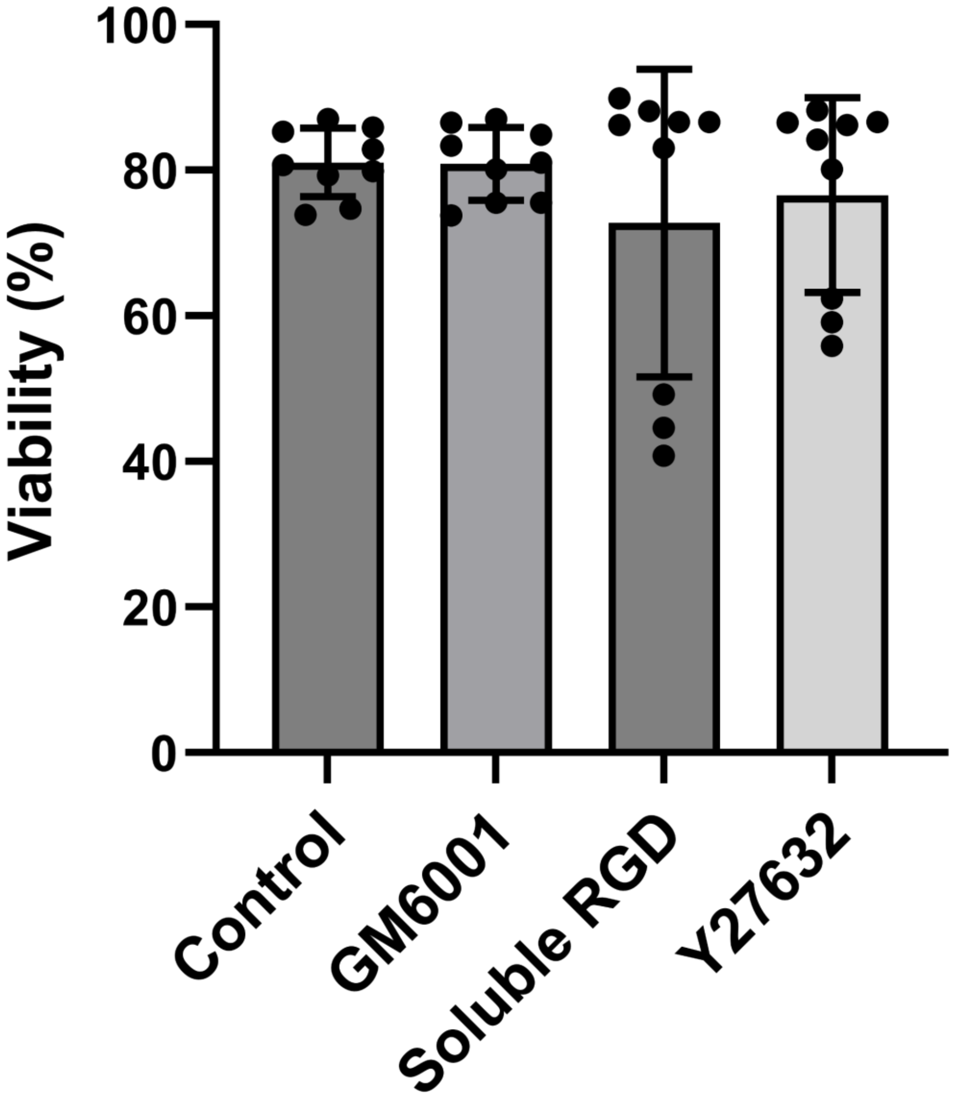

**Figure S2: Viability of primary NK cells after inhibitor treatment**. NK cells were treated with the inhibitors used in this study for 2 hours and the viability was determined by flow cytometry. Statistical significance was determined using a one-way ANOVA followed by a Tukey's multiple comparison test.

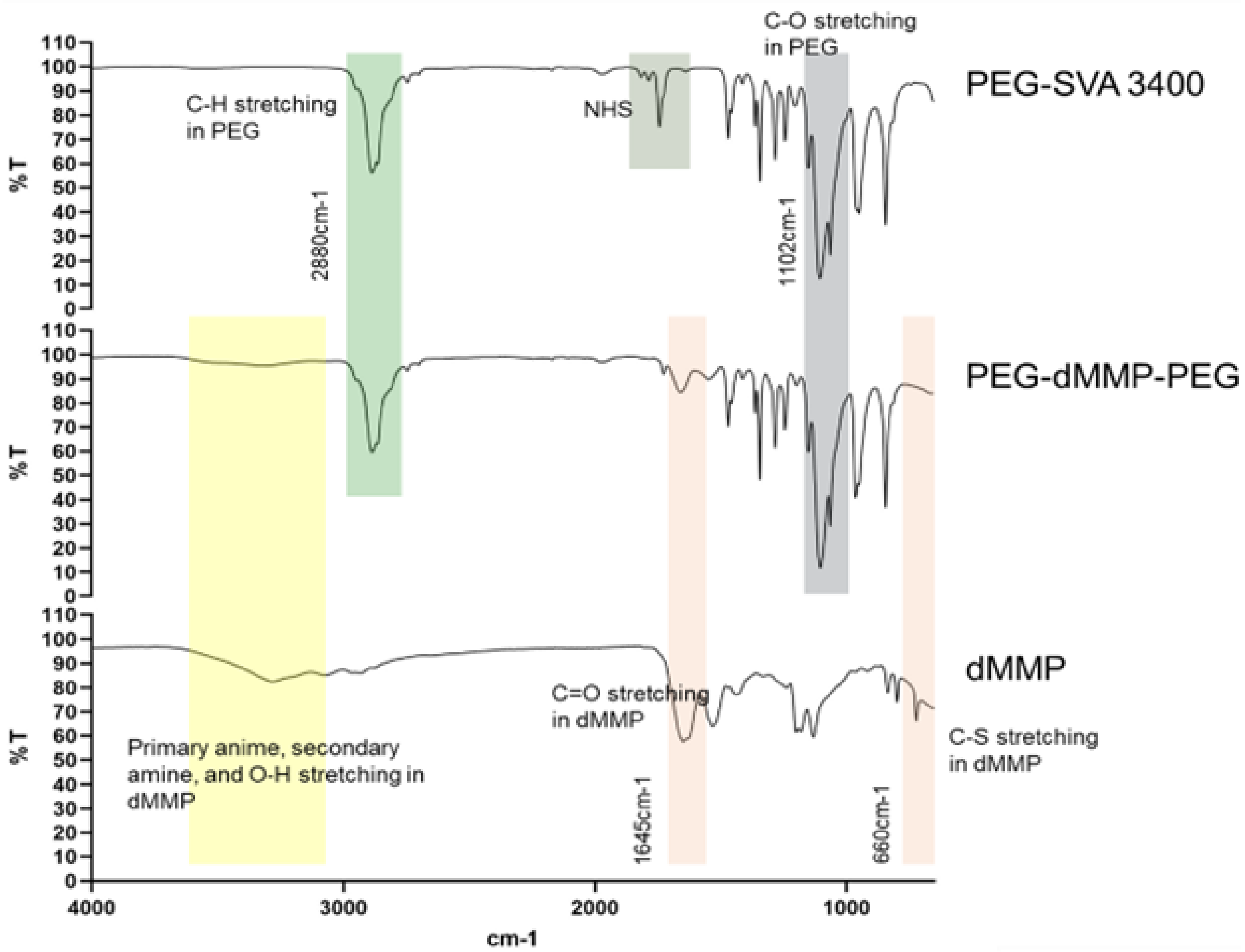

**Figure S3. FTIR spectrum to confirm functionalization of PEG with dMMP peptide.** Top is spectrum for acryl-PEG-SVA (MW 3400), middle is spectrum for PEG-dMMP-PEG, and bottom is spectrum for dMMP peptide.

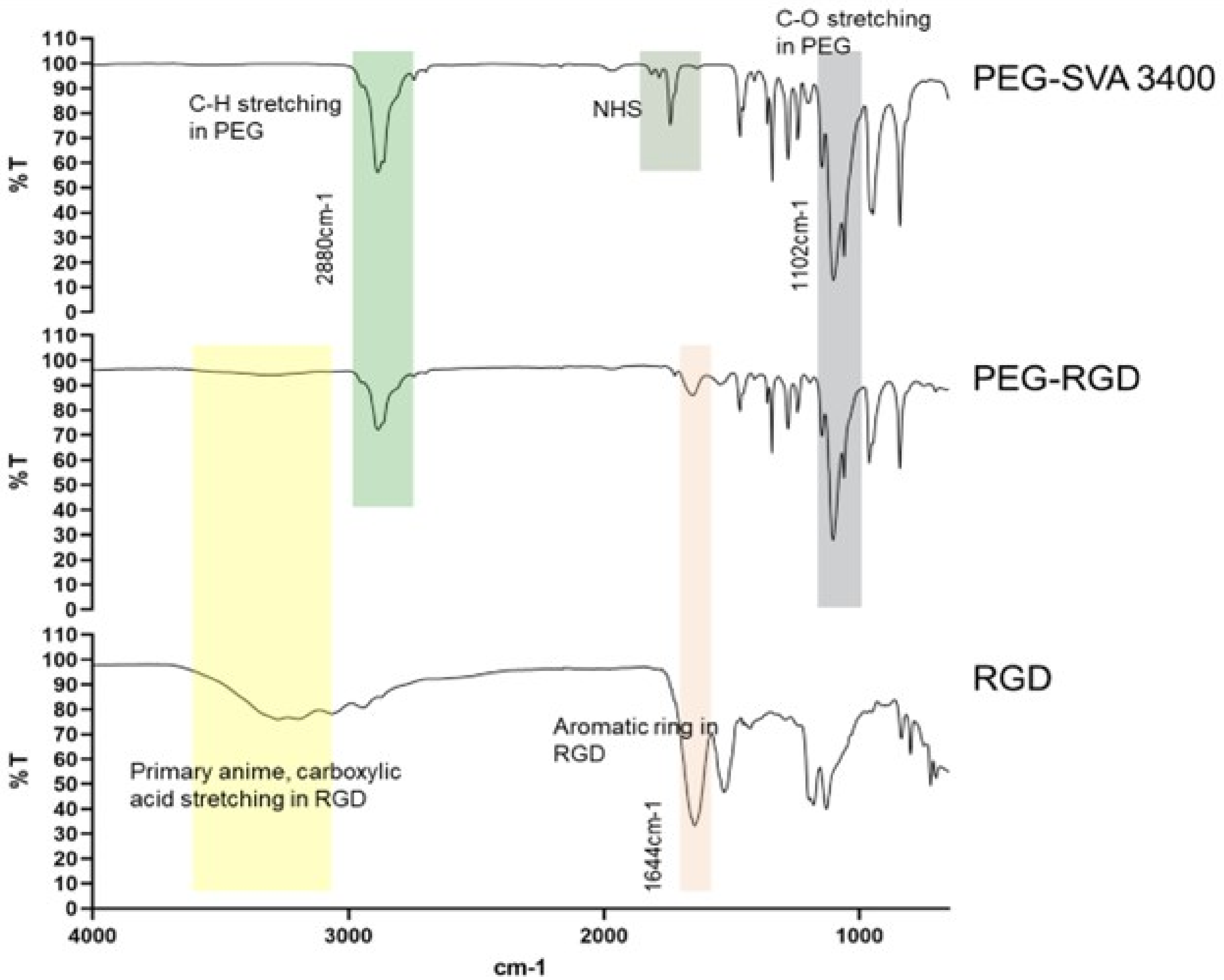

**Figure S4. FTIR spectrum to confirm functionalization of PEG with RGD peptide.** Top is spectrum for acryl-PEG-SVA (MW 3400), middle is spectrum for PEG-RGD, and bottom is spectrum for RGD peptide.

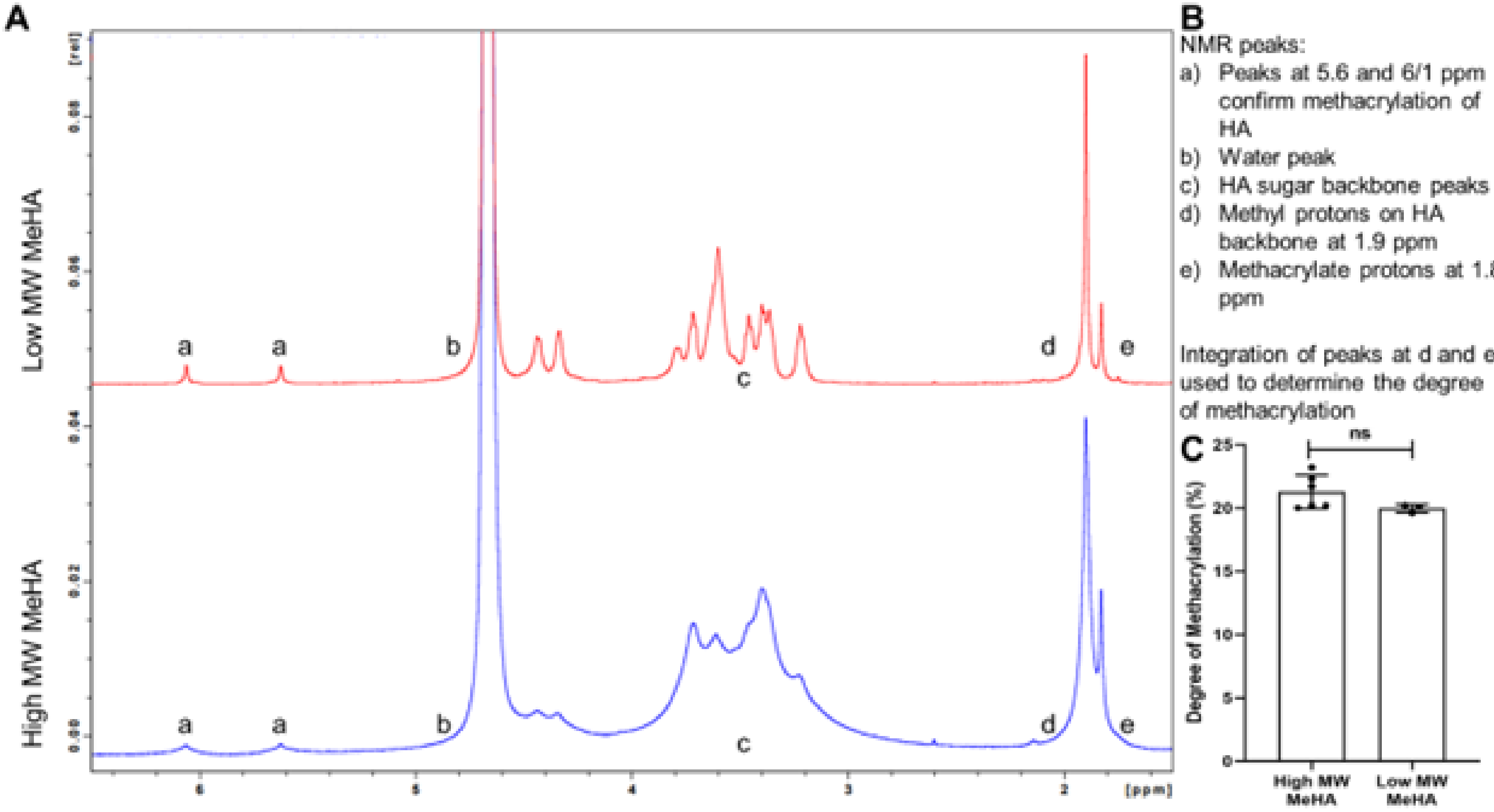

**Figure S5. HNMR analysis of LMW and HMW MeHA.** (A) Spectrums (pulse-acquired method) of LMW and HMW MeHA. (B) Key for the letter tags on the spectrums in A. (C) Quantification of the degree of methacrylation, based on the integration of peaks at d and e, for low and high MW MeHA.

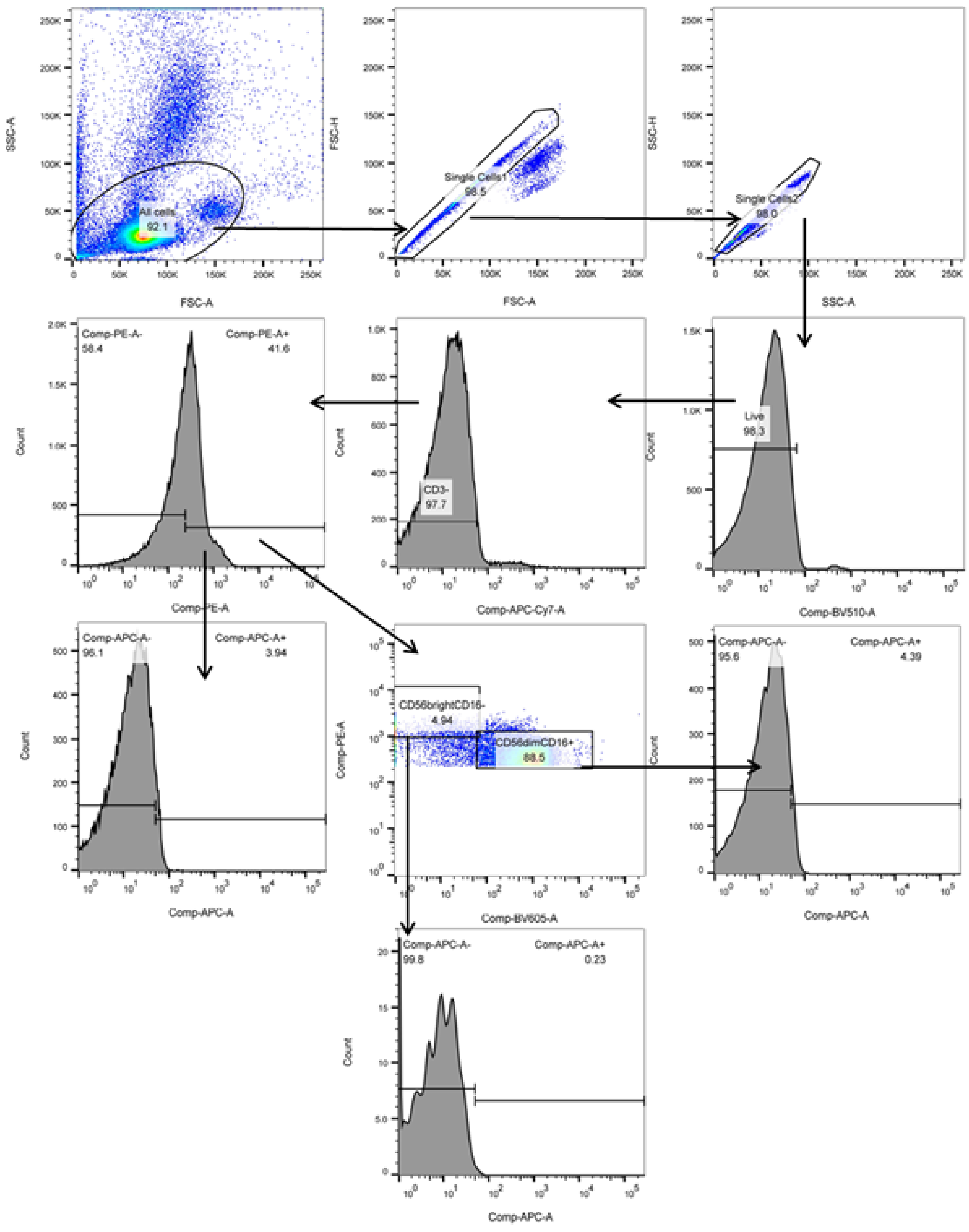

**Figure S6. Representative gaiting strategy for isolated primary NK cells.** Of note, only APC stain for CD3-CD56+, CD56brightCD16-, and CD56dimCD16+ is shown for simplification, but this was also for FITC, PerCP-Cy5.5, and BV421 for the other chemokine receptors.
